# Spectral Unmixing: A modular and reproducible Python package for directed and blind bleed-through correction in multidimensional microscopy stacks

**DOI:** 10.64898/2026.07.06.736825

**Authors:** Fabrizio Musacchio, Martin Fuhrmann

## Abstract

**Background:** Spectral bleed-through is a persistent source of bias in multi-channel fluorescence microscopy, where signal from one fluorophore is recorded in another detection channel. Routine correction workflows often remain fragmented across manual graphical procedures, laboratory-specific scripts, or method-specific blind-unmixing implementations with limited provenance.

**Results:** We present *spectral-unmixing*, an open-source Python package for reproducible directed and blind bleed-through correction in multidimensional microscopy stacks. The package combines directed two-channel correction with multiple coefficient-estimation strategies, optional bidirectional two-channel correction through explicit inversion of a two-by-two mixing model, and PICASSO-family blind unmixing for multichannel data. Input stacks from multiple microscopy file formats are normalized to a canonical axis order before processing, reducing the need for manual pre-formatting. Each processing run also writes a machine-readable sidecar file that records the effective configuration and estimated coefficients. Synthetic and real-data-derived benchmarks demonstrated robust correction across directed, bidirectional, and blind-unmixing settings. In fixed-alpha two-channel simulations, directed correction reduced target-channel normalized root mean squared error from approximately 0.029 to about 0.003.

In time-varying data, per-time-point estimation reduced mean absolute alpha error from approximately 0.099 to 0.003 compared with reference-time-point estimation, while bidirectional inverse-model correction reduced reciprocal-mixture channel errors from approximately 0.022–0.037 to about 0.004. Multichannel benchmarks further showed that blind-unmixing workflows can reduce residual inter-channel dependence while preserving fluorophore identity, and that source-sink priors provide a controllable alternative when plausible contamination pathways are known.

**Conclusions:** *spectral-unmixing*provides a modular, scriptable, and extensible platform that standardizes directed, bidirectional, and blind spectral-unmixing workflows, thereby lowering barriers to reproducible bleed-through correction in quantitative bioimage analysis.

## 1 Background

Multichannel fluorescence microscopy is routinely used to study spatial relationships and temporal interactions between labelled cellular structures in cell biology and neuroscience. However, when emission spectra overlap or detection windows are not cleanly separated, signal from one fluorophore can spill into the measurement channel of another fluorophore. This effect, often referred to as spectral bleed-through, crosstalk, or spillover, creates false-positive signal, distorts intensity-based readouts, and can bias the interpretation of apparent colocalization or structural dynamics[1–4]. In longitudinal imaging experiments, these errors can propagate further into time-resolved analyses, especially when image stacks include both time and z axes and when signal-to-noise ratios vary over the course of an acquisition. From a computational perspective, the problem is not only how to subtract one signal from another, but how to do so transparently and reproducibly. In everyday practice, spectral bleed-through correction is often handled by a mixture of manual brightness and contrast adjustments, spreadsheet-style note taking, GUI operations, macros, and laboratory-specific scripts. These approaches may produce visually acceptable results, yet they frequently leave no machine-readable provenance of which coefficient was used, how it was estimated, or whether different time points were treated differently. Similar reproducibility issues have been documented broadly for microscopy data handling and metadata interpretation[5–9].

At the same time, the existing software ecosystem is fragmented. Fiji/ImageJ provides a powerful and extensible platform for exploratory image analysis[10, 11], but unmixing workflows there are usually GUI-driven or plugin-specific rather than exposed through a dedicated, scriptable, provenance-oriented application programming interface (API). The original PICASSO formulation introduced a blind-unmixing criterion based on minimizing statistical dependence between channels without reference spectra[4], but that work primarily addressed a specific blind-unmixing scenario rather than a general software platform for routine directed, bidirectional, and multidimensional unmixing workflows. General hyperspectral-analysis packages, learning-based splitting methods, and microscope-vendor software each serve important purposes, but they are not primarily designed to provide a compact, open, provenance-oriented workflow for common fluorescence bleed-through correction in canonical microscopy stacks. In addition, microscopy file handling itself remains heterogeneous, with axis conventions and metadata interpretation varying across formats and libraries[5, 12]. Table 1 therefore summarizes the practical software gap addressed by *spectral-unmixing*. The comparison is deliberately pragmatic: it does not try to rank all image-analysis software, nor does it claim that general-purpose platforms cannot be adapted to a given task through custom macros or external scripts. Instead, it asks which capabilities are exposed directly and coherently for routine spectral-unmixing workflows relevant to microscopy users. The table does not suggest that existing platforms are inadequate. Fiji/ImageJ remains indispensable for exploratory image analysis, vendor-integrated suites remain central for acquisition-coupled microscopy workflows, and PICASSO established an influential blind-unmixing criterion. Rather, the comparison highlights that the available ecosystem is split across different methodological specializations. PySptools, HyperSpy, and HySUPP address general hyperspectral or multidimensional signal-analysis problems, but they are not centered on routine fluorescence bleed-through correction in canonical microscopy stacks[13–15]. PICASSO provides a strong blind-unmixing workflow, but it is not structured as a compact microscopy-stack package that also makes directed and bidirectional two-channel correction first-class workflows[4, 16]. Recent learning-based or content-aware approaches such as *µ*Split, denoiSplit, scSplit, and *λ*Split further demonstrate the breadth of current image-separation research: they emphasize trained image splitting, denoising, or content-aware spectral recovery, often with strong performance on their intended benchmarks. Their primary contribution, however, is a task-specific reconstruction model rather than a lightweight, method-agnostic workflow layer for routine bleed-through correction with interchangeable estimation strategies and explicit per-run provenance[17–20]. Taken together, these comparisons show that no single cited workflow simultaneously exposes directed two-channel correction, bidirectional inversion, blind multichannel unmixing, flexible multidimensional microscopy-stack handling, and readable per-run provenance as first-class features. This gap defines the design target of *spectral-unmixing*: a modular research-software package that standardizes how common linear bleed-through correction workflows are expressed, executed, documented, and shared.

**Table 1.** Comparison of software modes relevant to spectral bleed-through correction. The table emphasizes directly exposed workflow capabilities rather than every theoretically possible extension. “Partial” indicates that the capability can be approximated via manual steps, macros, or external code, but is not the central workflow offered by the tool. * marks open-source workflows. ^†^ marks proprietary workflows.

| Tool / workflow | Directed 2-ch | Bidirectional 2-ch | Blind multi-ch | TZCYX-aware scripting | Readable provenance |
| --- | --- | --- | --- | --- | --- |
| **GUI / macros** |  |  |  |  |  |
| Fiji / ImageJ* [10, 11] | Partial | No | Partial | Partial | No |
| **Python packages** |  |  |  |  |  |
| PySptools* [13] | No | No | No | No | No |
| HyperSpy* [14] | No | No | Partial | Partial | No |
| HySUPP* [15] | No | No | Yes | No | No |
| **Research workflows / method papers** |  |  |  |  |  |
| PICASSO* [4, 16] | Yes | Yes | Yes | No | No |
| μSplit* [17] | Partial | Partial | No | No | No |
| denoiSplit [18] | Partial | Partial | No | No | No |
| scSplit* [19] | Partial | Partial | No | No | No |
| λSplit [20] | Partial | No | Yes | No | No |
| **Proprietary software** |  |  |  |  |  |
| Vendor microscopy suites† [21–24] | Partial | Partial | Partial | Partial | Partial |
| **Our solution (Python package)** |  |  |  |  |  |
| *spectral-unmixing* * [25] | Yes | Yes | Yes | Yes | Yes |

Fiji / ImageJ refers to the public Fiji / ImageJ ecosystem as a general-purpose image-analysis platform rather than to any one specific spectral-unmixing plugin [10, 11]. PICASSO refers here to the open-source MATLAB / Python workflow family centered on the original blind-unmixing criterion and its released scriptable implementations [4, 16]. Vendor microscopy suites denotes representative proprietary software ecosystems such as ZEISS ZEN, Leica LAS X, Nikon NIS-Elements, and Evident / Olympus cellSens [21–24].

The specific role of *spectral-unmixing* is to provide a unified and extensible software layer for common linear spectral-unmixing tasks in microscopy: directed two-channel unmixing, bidirectional two-channel correction, and PICASSO-family blind unmixing. A central design decision is the use of OMIO as the input/output (I/O) layer[26], so that image stacks from multiple microscopy file formats enter the unmixing workflow in a canonical *TZCY X* representation with OME-compatible dimension semantics. This lets the package focus on mathematically explicit unmixing models and reproducible workflow behaviour rather than on format-specific parsing logic. These considerations lead directly to the design goals in Box 1. The goals are intentionally software-oriented because the intended contribution is broad utility and standardization: *spectral-unmixing* brings related unmixing workflows under one API, documents their assumptions, and records their effective settings in machine-readable form.

#### Box 1. Design goals of *spectral-unmixing*

*spectral-unmixing* was built around five explicit design goals:

1. **A small, discoverable core API.** The main workflow should remain centred on two entry points, unmix(…) and unmix picasso(…), so that routine directed and blind workflows stay scriptable and easy to learn.
2. **Explicit mathematical models and estimation logic.** Every unmixing path should correspond to a clearly documented linear model or optimization criterion, and coefficient provenance should remain explicit.
3. **Reproducibility by default.** Every output stack should be accompanied by a machine-readable sidecar report file recording the chosen method, effective parameters, and estimation details.
4. **OMIO-based multidimensional stack interoperability.** File-format heterogeneity should be delegated to OMIO so that downstream unmixing always operates on canonical *T ZCY X* stacks, including the common edge cases *T* = 1 and *Z* = 1.
5. **Modularity, extendibility, and workflow integration.** New alpha estimators, blind-unmixing routines, and optional helpers should be easy to add while keeping the package useful in Python-based bioimage-analysis pipelines and teaching material.

The main contribution of the package is therefore the creation of a modular, open, and reproducible software platform that unifies several commonly needed spectral-unmixing workflows under one interface. The package is distributed as free and open-source software under GPLv3, includes tests, continuous integration, tutorials, example datasets, and sidecar reports for every processed output[25, 27]. These design choices matter scientifically because they make spectral-unmixing analyses easier to audit, reuse, compare, and extend across microscopy projects, turning corrected image data into shareable and FAIR-oriented analysis artifacts rather than isolated outputs[7, 9].

## 2 Implementation

### 2.1 Software architecture

The implementation follows the design goals summarized in Box 1. To address point 1 therein, the package exposes a small public API with stable entry points and consistent parameter conventions across workflows. This is reflected in the separation between the lower user-level area and the central package workflow in Fig. 1: users provide an input image file, scientific settings, and scripted processing instructions, while the backend code remains unchanged. Directed two-channel workflows, including optional bidirectional correction, are exposed through unmix(…), whereas PICASSO-family blind-unmixing workflows are exposed through unmix picasso(…). Internally, these entry points dispatch to specialized components for coefficient estimation, blind-unmixing optimization, stack transformation, clipping, output writing, and reporting, but the user-facing interface remains compact and discoverable.

**Fig. 1.**
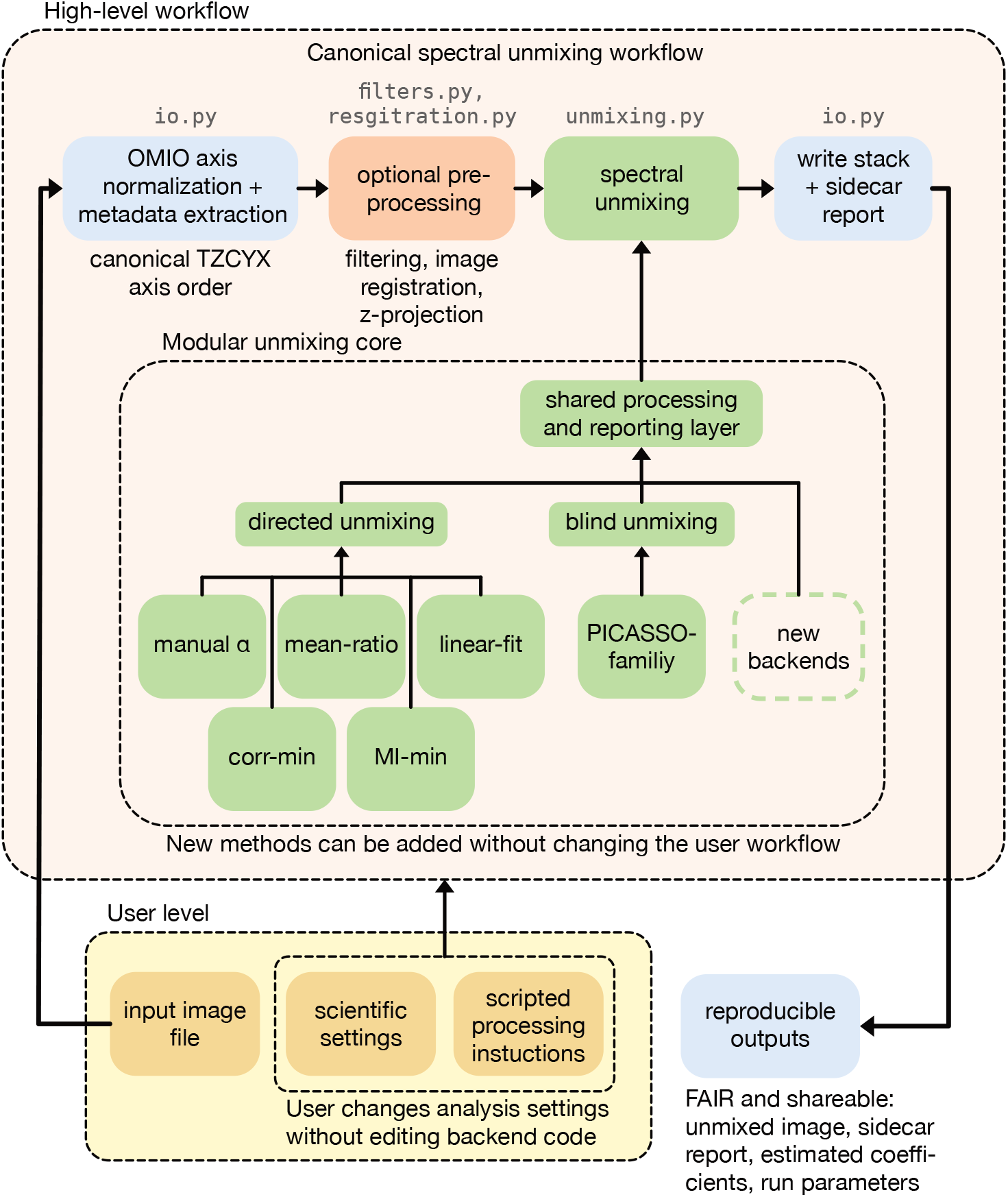
Software architecture and workflow logic of the *spectral-unmixing* package. The lower user-level area separates input image files, scientific settings, and scripted processing instructions from the backend implementation. The upper workflow band shows the canonical processing path: OMIO-based axis normalization and metadata extraction convert microscopy data into a canonical *T ZCY X* representation, optional processing helpers can be applied when needed, and the core spectral-unmixing routines operate on the normalized stack. The central modular unmixing core groups directed coefficient-based methods, PICASSO-family blind-unmixing backends, and future extensibility through a shared processing and reporting layer. The output path writes both the processed image stack and a sidecar report, allowing results and effective processing settings to be shared together.

To address point 4 of Box 1, microscopy file handling is treated as an explicit I/O boundary rather than as part of each unmixing method. In the upper workflow band of Fig. 1, this boundary is represented by OMIO-based axis normalization and metadata extraction before any unmixing-specific operation is applied[26, 28]. OMIO reads microscopy file formats and normalizes them to a canonical stack representation before the data enter the unmixing routines. Internally, the package therefore operates on microscopy stacks represented as

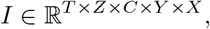

where *T* denotes time, *Z* axial planes, *C* channels, and *Y, X* spatial coordinates. This boundary is important practically: users can start from microscopy data as exported by their microscope software, while OMIO handles file-format parsing and axis normalization. Because the same canonical representation includes the common edge cases *T* = 1 and *Z* = 1, the downstream unmixing code can handle 2D two-channel images, single-time-point 3D stacks, and full 3D time-lapse stacks without shape-dependent branching in user code. This avoids making spectral-unmixing reproducibility depend on a laboratory-specific pre-conversion step.

To address point 5 of Box 1, the package separates core unmixing logic from workflow helpers. This is indicated by the central modular unmixing core in Fig. 1: directed estimators, PICASSO-family blind-unmixing backends, and future method backends connect through a shared processing and reporting layer. New alpha estimators can reuse the existing source-target preparation and mask interface. New blind-unmixing routines can reuse the same I/O, stack transformation, clipping, and reporting infrastructure. Optional filtering, projection, registration, and napari-viewer helpers are implemented outside the core unmixing routines so that they can support practical workflows without obscuring the mathematical assumptions of the unmixing methods themselves.

The same separation also defines how the package supports reproducible use. The user changes scientific settings and processing instructions at the script level, not by editing backend code. The output path then writes both the processed stack and its sidecar report. Thus, reproducibility is not treated as a separate post hoc documentation step, but as a consequence of the architecture: user-controlled settings enter the workflow explicitly, core methods process canonical stack data, and the resulting image and provenance information are emitted together.

### 2.2 Implemented mathematical framework

Point 2 of Box 1 is addressed by making the mathematical model behind each workflow explicit. The following subsections therefore define the linear models, coefficient-estimation rules, and dependence-minimization criteria implemented by the package.

They are included as part of the software implementation because the package is intentionally method-driven: each user-facing workflow corresponds to a model or optimization criterion that can be inspected, reproduced, and extended.

#### 2.2.1 Directed two-channel linear unmixing

##### Forward model

The basic directed two-channel model assumes that one measured channel is approximately a pure source channel and that the second measured channel contains both its own signal and a linear contribution from the source channel:

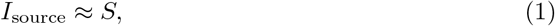

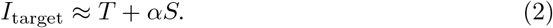

Here, *S* denotes the true source-channel signal, *T* the true target-channel signal, and *α* ≥ 0 the bleed-through coefficient from source into target. The corrected target channel is therefore

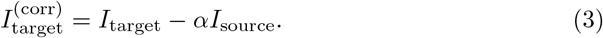

Optionally, negative output values are clipped to zero after subtraction because negative fluorescence intensities may be considered not physically meaningful in the final image:

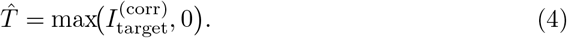

Only the target channel is modified; the source channel is preserved.

### Alpha modes: Where coefficients are obtained from

The package distinguishes between where an alpha value is obtained from and how it is estimated numerically. Let *X_t,z_*and *Y_t,z_* denote the source and target sub-volumes at time point *t* and z plane *z*. Then:

- **Fixed mode (**fixed) uses one user-supplied scalar *α* for all *t* and *z*.
- **Reference-time-point mode (**reference t) estimates one scalar *α*^ from all z-slices at a chosen reference time point *t*_ref_ and applies it to the full stack.
- **Per-time-point mode (**per t) estimates one scalar *α*^*_t_* for each time point from all z-slices at that time point.

Operationally, reference-time-point estimation assumes that the bleed-through factor is stable across time, whereas per-time-point estimation allows it to drift slowly over time. The latter is useful when global intensity levels change during an acquisition, but it may also absorb biologically meaningful time-dependent covariance if source and target signals genuinely co-vary.

### Optional preprocessing and alpha masks

For data-driven alpha estimation, the package can optionally apply a simple percentile-based preprocessing step to the source and target estimation volumes:

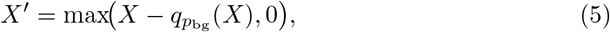

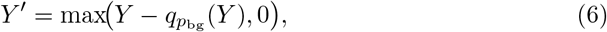

where *q_p_*_bg_ denotes the low-percentile intensity used as a rough background estimate. This preprocessing is used only for alpha estimation, not for the final image correction step in Eq. (3).

Alpha estimation is restricted to a mask of bright source voxels:

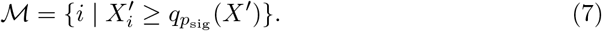

Optionally, the mask can be further constrained to comparatively low target intensities,

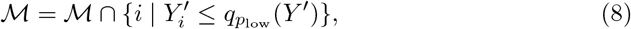

which is helpful when the user wants to emphasize voxels likely dominated by bleed-through rather than by genuine target signal. If such a compound mask becomes too small, the implementation falls back to the source-only mask and records that fallback in the sidecar report.

### Alpha estimation methods

Given prepared source and target vectors (*X_i_’*,*Y_i_’*) on the estimation mask *M*, the package implements five estimation methods.

#### Manual estimator *(manual)*

The user provides *α* explicitly. This mode is scientifically preferable when a coefficient has been measured on an appropriate single-label control recorded under matching acquisition settings.

#### Mean-ratio estimator *(mean ratio)*

The default data-driven method computes

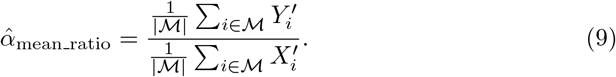

This is simple, stable, and intentionally close to the intuitive “average bleed-through fraction” interpretation.

#### Linear-fit estimator (linear fit)

The masked least-squares fit without intercept is

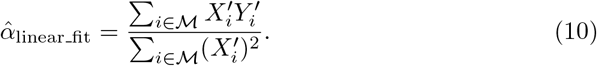

Because the background has already been handled separately, no intercept is fitted.

***Correlation-minimization estimator* (**corr min).

For correlation minimization, the corrected target is defined as *Y* ^′^ — *αX*^′^, and *α* is chosen by bounded scalar optimization on [0*, α*_max_]:

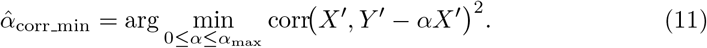

This explicitly minimizes residual linear dependence between source and corrected target.

#### Mutual-information minimization *(mi min)*

Mutual-information minimization uses a two-channel PICASSO-like criterion:

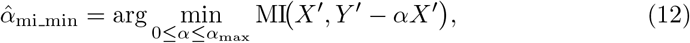

where MI is estimated from a two-dimensional intensity histogram. This method is especially attractive when the residual source-target relationship is not adequately described by correlation alone, although it may be computationally heavier and more variable than the simpler ratio-based estimators.

#### Bidirectional two-channel unmixing

Some imaging setups exhibit bleed-through in both directions. In that case the package can solve a 2 × 2 linear mixing system instead of applying two sequential subtractions:

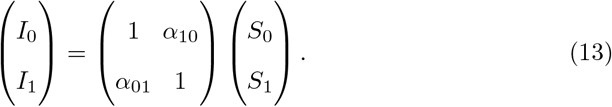

Provided that 1 — *α*_01_*α*_10_ /= 0, the unmixed signals follow by matrix inversion:

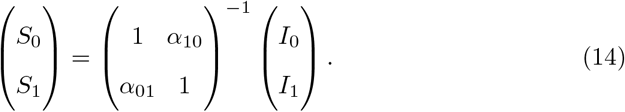

This formulation is preferable to sequential subtraction because sequential order would otherwise influence the result.

#### 2.2.2 PICASSO-family blind unmixing

##### General model

For multi-channel blind unmixing, the package follows the linear mixing viewpoint of the original PICASSO publication[4]. Let *I* ∈ R*^C^*^×^*^N^* denote *C* measured channels flattened over all spatial samples and *F* ∈ R*^C^*^×^*^N^* the unknown fluorophore signals, under the assumption that the number of measured channels equals the number of fluorophores:

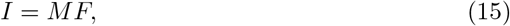

where *M* is an unknown mixing matrix. The goal is to estimate an unmixing transform *U* such that

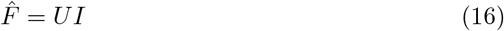

shows reduced statistical dependence between channels.

##### **Matlab_3c**: close Python port of the published three-channel workflow

The matlab 3c implementation aims to reproduce PICASSO’s original three-channel MATLAB logic as closely as possible. Let *F* ^(*k*)^ ∈ R^3×^*^N^* denote the current three-channel estimate at iteration *k* after per-channel maximum normalization, low-percentile background removal, and slice-wise pixel binning. For each ordered sink-source pair (*i, j*) with *i* /= *j*, the workflow estimates a subtraction coefficient by minimizing histogram-based mutual information:

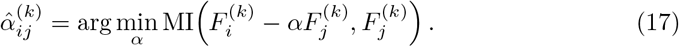

The resulting pairwise coefficients are clipped and inserted into an incremental update matrix

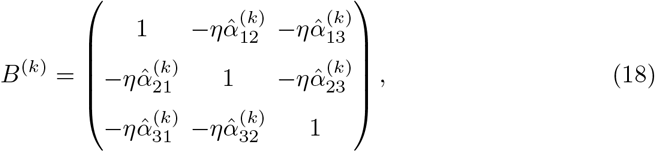

where *η* is the step size. In the package implementation, this update is then applied in the same row-wise sequential order as in the released MATLAB workflow, so that later rows in the same sweep can already depend on channels updated earlier in that sweep. Positivity is enforced intermittently by clipping negative intermediate values to zero.

##### Matlab_n: explicit package-level generalization to N channels

The PICASSO publication formulates blind unmixing for an arbitrary number of fluorophores, whereas the released reference MATLAB workflow operates in a three-channel setting. In *spectral-unmixing*, the same released iterative update logic is generalized to *N* channels through matlab n. Concretely, for each ordered channel pair (*i, j*) with *i* /= *j*, a mutual-information-minimizing subtraction coefficient is estimated and applied sequentially as

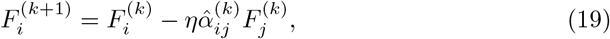

with the corresponding row update carried through the running unmixing matrix. Equivalently, one may view the method as constructing an *N* × *N* analogue of *B*^(*k*)^ whose diagonal entries remain one and whose off-diagonal entries store the pairwise subtraction terms 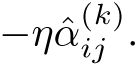 The preprocessing, histogram-based mutual-information criterion, and sequential incremental updates are retained; only the channel iteration is generalized. Accordingly, matlab n should be understood as a package-level generalization of the original three-channel workflow rather than as a new blind-unmixing criterion.

#### source sink n: explicit source-sink modeling

The package additionally provides source sink n, a source-sink formulation intended for scenarios where users already have some intuition about which channels act mainly as sinks and which channels are plausible sources. Let *S* ∈ {—1, 0, 1}*^C^*^×^*^C^* encode allowed interactions, with *S_jj_*= 1, *S_ij_*= —1 if channel *i* is allowed to contribute to sink channel *j*, and *S_ij_* = 0 otherwise. After global intensity normalization and low-percentile background handling, the corrected sink is modeled as

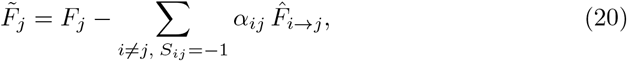

with prepared source terms

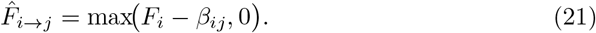

Here, *α_ij_* is the source-to-sink subtraction coefficient and *β_ij_* is an optional small background offset associated with that source-sink relation. By default, the implementation optimizes all coefficients contributing to one sink jointly by minimizing the summed histogram-based mutual information

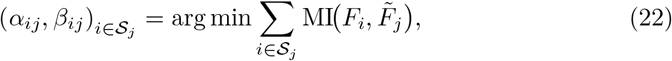

subject to 0 ≤ *α_ij_* ≤ *α*_max_ and, when background optimization is enabled, 0 ≤ *β_ij_* ≤ *β*_max_. A greedier sequential fallback remains available in the API, but the joint optimization is the default because it more closely matches the source-sink viewpoint implemented in the napari PICASSO plugin[16]. This formulation is particularly useful when prior knowledge from fluorophore selection, filter layout, or detector design suggests that only specific channels are plausible contamination sources for a given sink channel. In that regime, the source-sink graph acts as an explicit directional prior: it can prevent biologically or instrumentally implausible subtraction pathways, reduce accidental signal removal from channels that should primarily remain sources, and make the correction behaviour easier to control than in a fully unrestricted blind-unmixing workflow.

Our implementation is intentionally not a one-to-one port of the original plugin. The latter optimizes its source-sink objective with a neural mutual-information estimator, whereas *spectral-unmixing* uses a histogram-based mutual-information objective together with deterministic numerical optimization and optional joint background fitting. We decided against a neural estimator for two reasons. First, the histogram-based criterion is simpler to understand and easier to reproduce across platforms. Second, the neural estimator would introduce a PyTorch dependency that is not otherwise needed for the rest of the package. Thus, we intentionally chose a simpler, more portable, and computationally lighter implementation that still preserves the source-sink modeling idea.

### 2.3 Reproducibility and output reporting

Point 3 of Box 1 is addressed at the output layer. As shown by the right-hand output path in Fig. 1, each processing run writes two linked artifacts: the processed image stack and an adjacent machine-readable sidecar report. These artifacts are generated by the workflow itself rather than by optional user note taking. They therefore provide a compact audit trail for downstream analyses, manuscript figures, shared example datasets, and re-analysis by other users.

The first artifact is the unmixed image stack. It is written through OMIO in an OMIO-supported microscopy stack format and preserves the intended multidimensional stack semantics used internally by the package. This makes the processed data usable outside the original script: the image can be reopened for visualization, used in downstream quantitative analyses, shared with collaborators, or deposited together with processed results. The second artifact is the sidecar report written next to the processed stack. It records the input and output file paths, stack shape and axis order, selected workflow, channel configuration, effective user parameters, estimated coefficients or unmixing matrices, relevant optimization settings, and warnings or fallbacks when applicable.

This output design links the user-level and backend-level parts of the architecture. The user-level settings shown in the lower part of Fig. 1 are not merely transient script states. They are propagated into the sidecar report together with the method-specific results generated by the backend. As a result, a processed image can be interpreted together with the settings that produced it, and another user can reconstruct the relevant analysis choices without access to a graphical session or undocumented laboratory notes.

The reporting strategy therefore supports transparency, shareability, and FAIR reuse. A corrected image can be inspected later with explicit knowledge of the channel mapping, coefficient-estimation strategy, estimated parameters, and output settings, rather than relying on manual reconstruction of the workflow. In this sense, the sidecar report is not only a convenience feature but a structural part of the workflow: together with the processed stack, it turns a spectral-unmixing result into a documented and transferable analysis artifact.

### 3 Benchmark design and evaluation metrics

To evaluate the package quantitatively, we generated four families of synthetic benchmarks and complemented them with three real-data-based microscopy examples. The benchmark construction and evaluation metrics define the analyses reported in the Results section.

#### Fixed-alpha two-channel benchmark

We generated 10 synthetic *T* × *Z* × *Y* × *X* = 4 × 3 × 96 × 96 source and target stacks from Gaussian blobs with mild spatial overlap, added Gaussian noise (*σ* = 0.6), and mixed the target channel with a fixed ground-truth bleed-through coefficient *α* = 0.28. The methods manual, mean ratio, linear fit, corr min, and mi min were evaluated in reference t mode.

#### Time-varying two-channel benchmark

To test reference t versus per t, we generated 10 synthetic stacks with *T* = 6 and let the ground-truth coefficient vary linearly from 0.18 to 0.38 across time while keeping the spatial signal model similar to the fixed-alpha benchmark.

#### Bidirectional two-channel benchmark

To evaluate reciprocal bleed-through correction, we generated 10 synthetic stacks with two true channels and mixed them in both directions according to

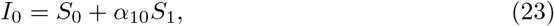

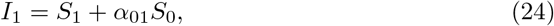

using *α*_01_ = 0.32 and *α*_10_ = 0.18. The same five alpha-estimation methods were then applied independently to the forward and reverse directions, and the resulting coefficients were used in the package’s bidirectional 2 × 2 inverse model. In this reciprocal setting, non-mean-ratio estimators were evaluated with a less restrictive source-mask threshold (signal percentile = 50 rather than 99) because the bidirectional mixtures benefited from retaining more source voxels during coefficient estimation.

#### Blind-unmixing benchmark

For 3-channel and 5-channel settings, we generated 5 sparse fluorescence scenes per setting, mixed them with known full-rank mixing matrices, added Gaussian noise (*σ* = 0.2), and evaluated matlab 3c, matlab n, and source sink n as applicable.

#### Metrics

The benchmarks use three families of quantitative metrics: reconstruction metrics for directed two-channel correction, coefficient-error metrics for bidirectional correction, and dependence-versus-recovery metrics for blind unmixing.

For directed two-channel benchmarks, the primary reconstruction metric is the normalized root mean squared error (NRMSE) between the corrected target image *T*^^^ and the ground-truth target image *T*,

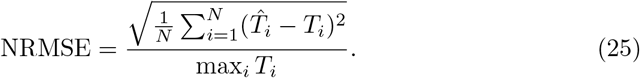

This quantity summarizes how accurately the target channel is recovered after correction, normalized by the target intensity scale. In addition, we report the residual Pearson correlation between the measured source channel and the corrected target channel. This residual correlation is not used as the main reconstruction metric, but it is informative as a compact measure of how much source-like structure remains in the corrected target after subtraction.

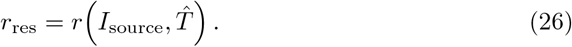

Here, *r*_res_ is a signed correlation for one specific source-target pair. It is therefore closely related to the correlation-based dependence measures used later for blind unmixing, but it is not the same metric: in the directed two-channel setting there is only one relevant pair, and the sign remains informative because slightly negative values can indicate over-compensation after subtraction.

For the two-channel biological example, no ground-truth target image is available. In that case, we additionally report the fraction of target-channel pixels that would become negative before the final non-negativity clipping step,

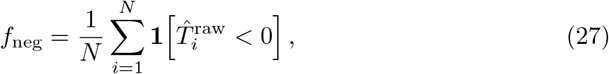

where 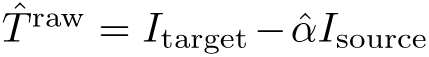 denotes the raw subtraction result prior to clipping. We use *f*_neg_ as a practical proxy for subtraction aggressiveness: larger values indicate that a greater fraction of the target image would need to be rescued by clipping and therefore suggest a higher risk of over-subtraction.

For bidirectional benchmarks, we analogously report channel-specific NRMSE values for both recovered channels, together with the mean absolute error across the two estimated bleed-through coefficients. This combination is useful because reciprocal mixtures require both accurate image recovery and accurate estimation of two coupled mixing parameters. For the five-channel simulation example derived from real microscopy data, channel-wise ground truth is available, so the blind-unmixing dependence and recovery metrics defined below can be evaluated directly.

For blind-unmixing benchmarks, the evaluation must separate two related but non-identical goals: reducing statistical dependence between output channels, and recovering the underlying fluorophore channels faithfully. To quantify residual channel coupling within the recovered stack itself, we report a pairwise correlation-sum metric. For recovered channels *U*_1_*,…,U_C_*, flattened over pixels or voxels, we define

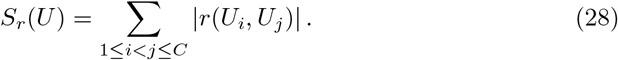

In other words, we consider all unordered channel pairs (*i, j*), compute the Pearson correlation for each pair, take the absolute value, and sum over all pairs. The absolute value prevents positive and negative correlations from canceling each other. A lower value of *S_r_*(*U*) therefore means that the recovered channels have become more decorrelated from one another. For a stack with *C* channels, this sum contains 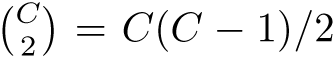 terms, so it is not bounded above by *C* and should not be interpreted as a per-channel recovery score. In the special case *C* = 2, the same basic Pearson-correlation concept is being measured as in *r*_res_, but the role of the metric differs: *r*_res_ is a signed one-pair diagnostic for directed source-target correction, whereas *S_r_*(*U*) is an absolute multi-pair aggregate dependence score for blind unmixing.

The corresponding pairwise mutual-information score is defined analogously,

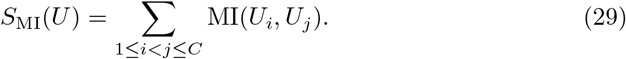

This metric again quantifies residual dependence within the recovered output channels, but now in a more general statistical sense. Whereas Pearson correlation primarily captures linear dependence, mutual information can remain positive even when linear correlation is weak, and therefore helps detect residual nonlinear coupling between channels. For both *S_r_*(*U*) and *S*_MI_(*U*), lower values indicate better channel separation in the sense of reduced residual inter-channel dependence.

To complement these dependence scores, we also report a best-match fluorophore-recovery score relative to the known ground-truth fluorophore channels. Let *G*_1_*,…,G_C_* denote the ground-truth channels and *U*_1_*,…,U_C_* the recovered channels. Because blind unmixing can involve both scale ambiguity and channel-permutation ambiguity, we do not assume that recovered channel index *i* must correspond directly to ground-truth channel index *i*. Instead, for each ground-truth channel we consider the strongest matching recovered channel, for example via the absolute Pearson correlation

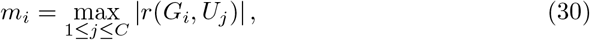

and summarize these matches through

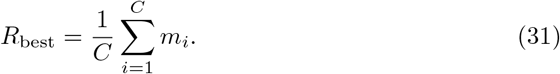

In the implementation, the actual assignment is obtained with the Hungarian algorithm on the matrix of absolute Pearson correlations so that each recovered channel is matched consistently and only once. Conceptually, however, *R*_best_ asks how well each true fluorophore channel can be recovered somewhere in the output, irrespective of output order. Higher values therefore indicate better fluorophore recovery. This distinction is important for interpreting the benchmarks: lower residual dependence does not automatically imply better recovery of the underlying fluorophore channels, and the Results section shows concrete cases in which these two objectives trade off against each other.

## 4 Results

### 4.1 Directed two-channel unmixing recovers fixed bleed-through coefficients accurately

On the fixed-alpha synthetic benchmark, all correction methods substantially improved the target reconstruction relative to the mixed baseline (Fig. 2, Table 2). The representative image panels in Fig. 2a show the source signal, the ground-truth target, the mixed target, and one corrected target realization. Without correction, the mixed target channel had an NRMSE of 0.0294 ± 0.0033 and a source-target correlation of 0.194 ± 0.021. In contrast, using the ground-truth manually supplied coefficient reduced the NRMSE to 0.0031 ± 0.0003 and shifted the residual correlation slightly negative (—0.076 ± 0.006), consistent with mild overcompensation from clipping and noise at the benchmark’s operating point.

**Fig. 2.**
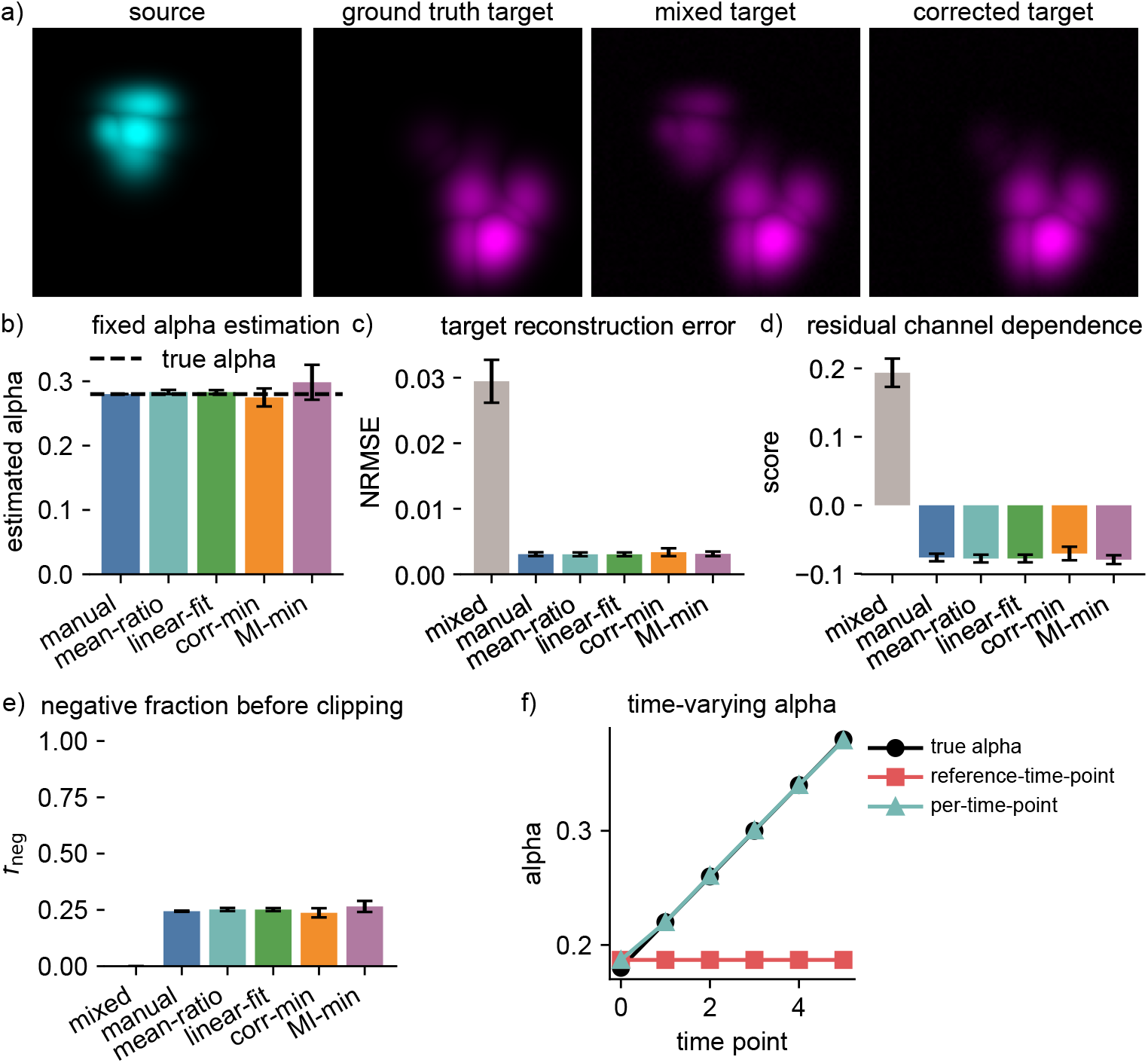
Directed two-channel benchmark. Panel a) shows a representative synthetic scene with source signal, ground-truth target signal, mixed target signal, and the target corrected by the default mean-ratio estimator. Panel b) to e) summarize coefficient recovery, reconstruction quality, residual dependence, and pre-clipping negative-fraction behaviour across methods, respectively. All data-driven estimators recover the fixed bleed-through coefficient closely on the controlled benchmark, and all markedly improve target reconstruction relative to the mixed baseline. Panel f) shows the additional time-varying-alpha benchmark, in which per-time-point estimation follows the true drift whereas reference-time-point estimation remains fixed.

**Table 2.** Summary of the fixed-alpha directed two-channel benchmark. Values are mean *±* standard deviation across 10 synthetic scenes.

| Method | Estimated $\alpha$ | NRMSE | Residual correlation |
| --- | --- | --- | --- |
| Mixed baseline | – | $0.0294 \pm 0.0033$ | $0.194 \pm 0.021$ |
| Manual (true $\alpha$) | $0.280 \pm 0.000$ | $0.0031 \pm 0.0003$ | $-0.076 \pm 0.006$ |
| Mean-ratio | $0.283 \pm 0.003$ | $0.0030 \pm 0.0003$ | $-0.078 \pm 0.006$ |
| Linear-fit | $0.283 \pm 0.003$ | $0.0030 \pm 0.0003$ | $-0.078 \pm 0.006$ |
| Corr. minimization | $0.275 \pm 0.014$ | $0.0034 \pm 0.0006$ | $-0.070 \pm 0.010$ |
| MI minimization | $0.298 \pm 0.027$ | $0.0031 \pm 0.0003$ | $-0.079 \pm 0.006$ |

Among the data-driven estimators, the mean-ratio and linear-fit approaches performed almost identically, both estimating *α* ≈ 0.283 and reducing NRMSE to approximately 0.0030. This close agreement is visible in the coefficient-recovery panel in Fig. 2b and the reconstruction-error panel in Fig. 2c. The optimization-based criteria also performed well but were more variable: correlation minimization yielded 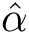 = 0.275±0.014, whereas mutual-information minimization yielded 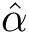 = 0.298±0.027. Residual dependence after correction (Fig. 2d) was similarly low for all corrected methods. The additional negative-fraction metric in Fig. 2e showed only modest differences in subtraction aggressiveness across methods in this controlled synthetic setting. The fraction of target pixels that would have become negative before clipping was 0.244 ± 0.003 for the manual-oracle correction, 0.252 ± 0.007 for mean-ratio, 0.251±0.007 for linear-fit, 0.237±0.020 for correlation minimization, and 0.265±0.025 for mutual-information minimization. Thus, in contrast to the more heterogeneous biological example analyzed later in this section, the synthetic benchmark indicates that the different estimators achieve very similar reconstruction quality with only comparatively small differences in clipping demand. These results are consistent with the intended role of the package: it exposes multiple scientifically interpretable estimation strategies rather than imposing one universally optimal estimator.

### 4.2 Per-time-point alpha estimation is advantageous in time-varying data

The time-varying benchmark shown in Fig. 2f revealed a clear difference between reference-time-point and per-time-point alpha estimation. When the true coefficient drifted from 0.18 to 0.38, reference-time-point estimation naturally remained locked to the initial estimate, yielding a mean absolute alpha error of 0.0988 ± 0.0015 and an NRMSE of 0.0124 ± 0.0013. In contrast, per-time-point estimation tracked the time-dependent coefficient closely, reducing the mean absolute alpha error to 0.0028±0.0018 and the NRMSE to 0.0030 ± 0.0002. This behaviour matters in practice because it shows that the choice of alpha mode directly changes correction quality whenever bleed-through or acquisition conditions drift over time.

### 4.3 Bidirectional inverse-model correction resolves reciprocal bleed-through

The bidirectional benchmark shows that reciprocal bleed-through is a meaningfully harder estimation problem than one-direction correction, because each measured channel is itself contaminated by the other. The three image rows in Fig. 3a illustrate this progression from ground truth to mixed observation to bidirectionally corrected reconstruction. In the mixed baseline, channel-specific NRMSE values were 0.0224 ± 0.0033 for channel 0 and 0.0372 ± 0.0055 for channel 1. Using the ground-truth bidirectional coefficients in the explicit 2 × 2 inverse model reduced these to 0.0036 ± 0.0005 and 0.0036 ± 0.0006, respectively, while lowering the measured inter-channel correlation from 0.571 ± 0.020 to 0.146 ± 0.031 and the corresponding mutual-information score from 0.431 ± 0.027 to 0.029 ± 0.009 (Fig. 3).

**Fig. 3.**
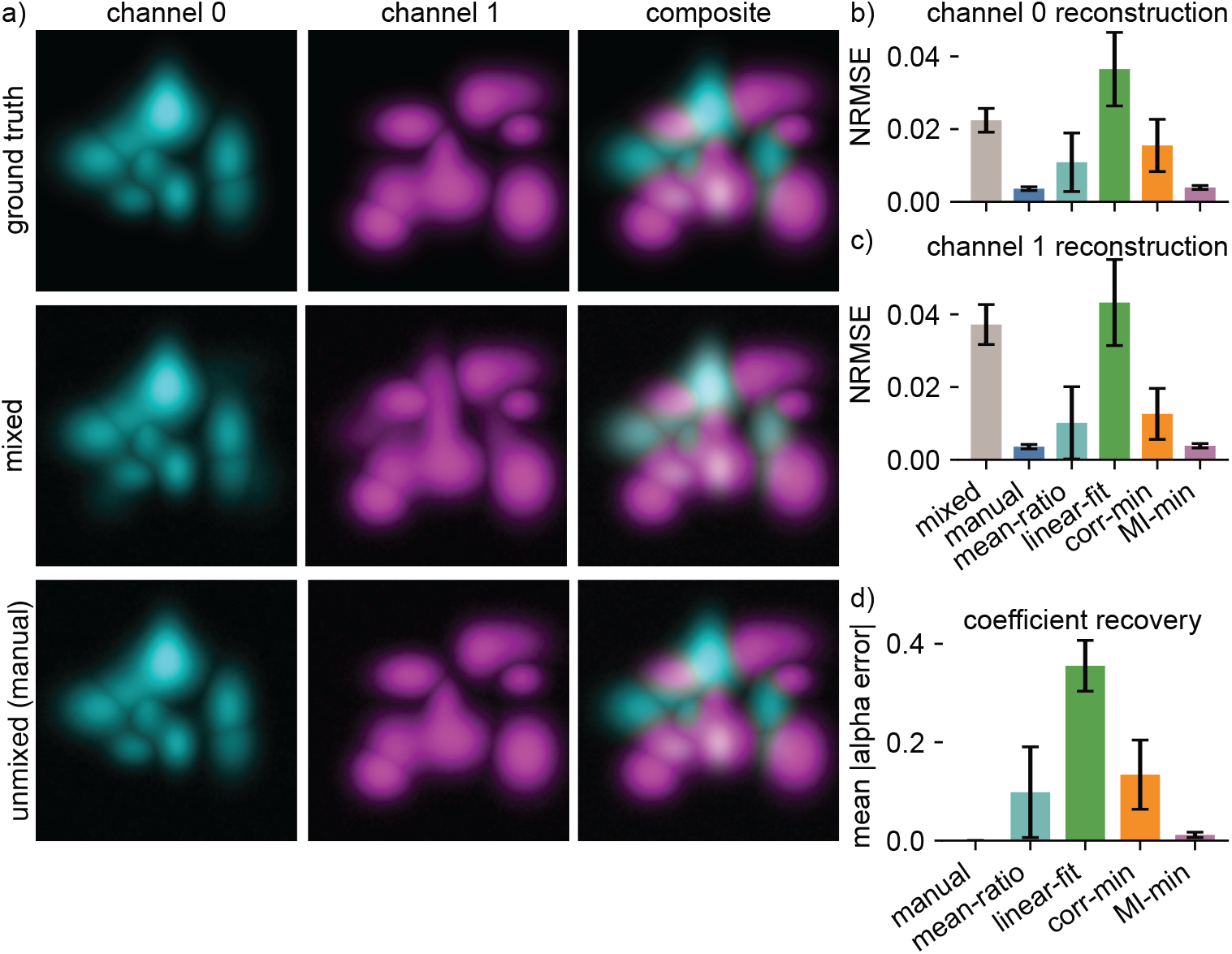
Bidirectional two-channel benchmark. The top row of panel a) shows a representative synthetic reciprocal-mixture scene as ground truth (from left to right: channel 0, channel 1, and composite). The middle row of panel a) shows spectrally mixed measurements of the same ground truth, and the lower row shows the bidirectionally unmixed result obtained with the true manual coefficients. Panel b) to d) summarize channel-specific reconstruction errors and bidirectional coefficient recovery across estimation methods. Reciprocal mixing is substantially harder than the one-direction case, but the explicit 2 *×* 2 inverse model still recovers both channels well when coefficients are accurate.

Among the data-driven estimators, the mutual-information minimization criterion performed best overall on this benchmark and came close to the manual-oracle solution, yielding channel-specific NRMSE values of 0.00395 ± 0.00054 and 0.00385 ± 0.00061 together with a mean absolute coefficient error of 0.0117±0.0054. The channel-specific reconstruction panels in Fig. 3b and c, together with the coefficient-recovery panel in Fig. 3d, make the same ranking visible graphically. Mean-ratio estimation remained usable but still overestimated the reciprocal coefficients, with mean estimates of *α*^_01_ ≈ 0.41 and *α*^_10_ ≈ 0.28. Correlation minimization showed intermediate performance, whereas linear-fit estimation was the least stable bidirectional estimator under this reciprocal-mixture setting, strongly overestimating both coefficients. Taken together, these results indicate that the package’s bidirectional mode is useful and well justified, but that reciprocal mixtures amplify the difficulty of data-driven coefficient estimation relative to the simpler one-direction setting.

### 4.4 PICASSO-family workflows reveal a trade-off between dependence reduction and fluorophore fidelity

On the blind-unmixing benchmarks, the MATLAB-style PICASSO workflows consistently reduced inter-channel dependence while preserving very high correspondence to the underlying fluorophore channels (Figs. 4 and 5). The representative image grids in Fig. 4a and Fig. 5a show that the MATLAB-style reconstructions remain visually close to the ground-truth channel organization, whereas the source-sink reconstructions appear more aggressively subtracted. In 3-channel scenes, the pairwise correlation sum dropped from 1.113 ± 0.308 in the mixed data to 0.469 ± 0.245 for both the MATLAB-style 3-channel and MATLAB-style *N*-channel workflows. The corresponding mutual-information sum decreased from 1.253 ± 0.147 to 0.470 ± 0.076, while best-match recovery correlations remained at 0.9961 ± 0.0007. The per-channel GT-correlation panels (Fig. 4e and Fig. 5e) show the same pattern directly at channel level: same-index correlations remain close to unity for the MATLAB-style workflows across the representative channels.

**Fig. 4.**
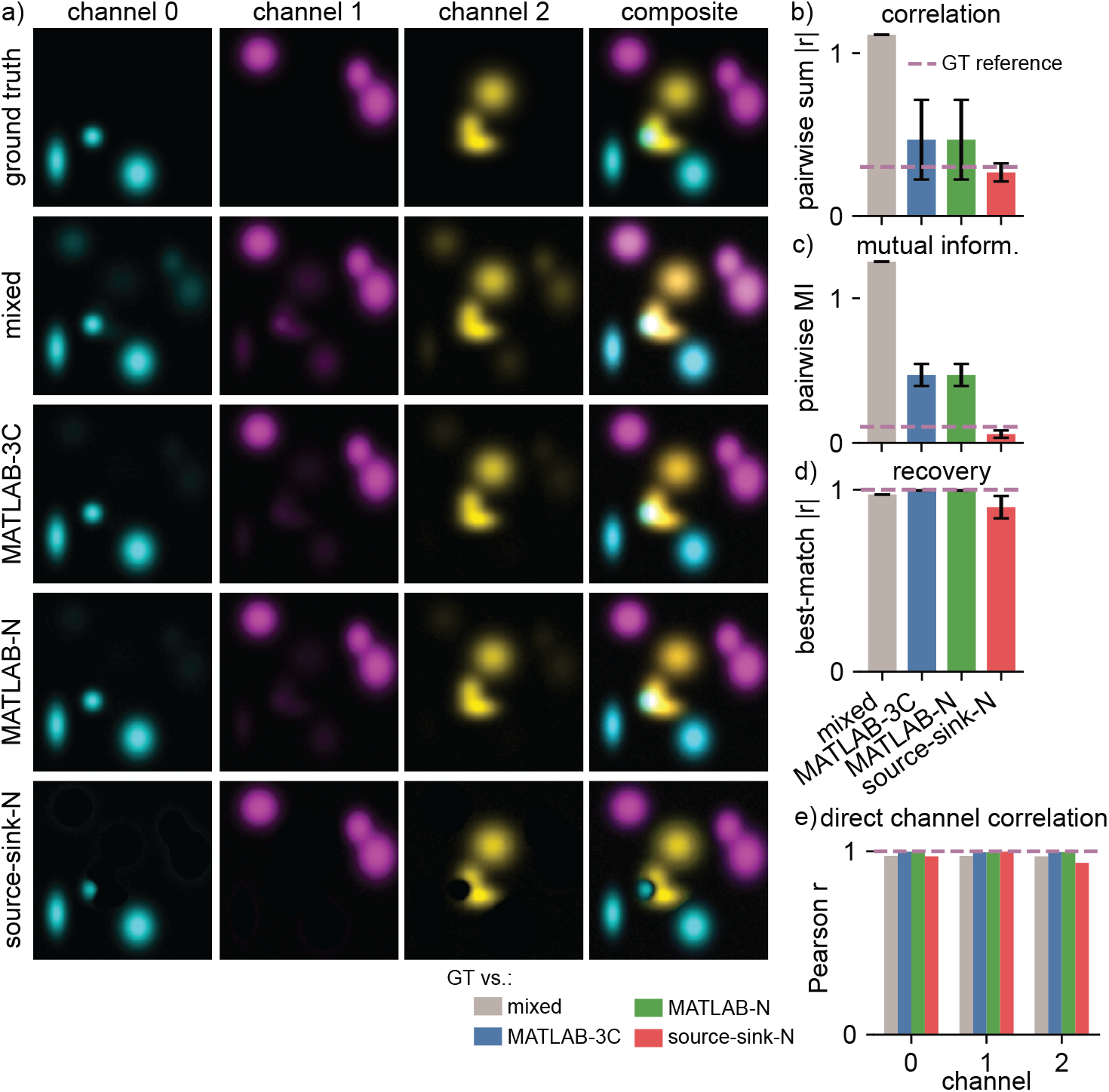
PICASSO-family blind-unmixing benchmark: 3-channel case. Panel a) shows individual panels of the ground-truth fluorophore channels, the spectrally mixed measurements, and the unmixed outputs obtained by the MATLAB-style 3-channel, MATLAB-style *N*-channel, and source-sink *N*-channel workflows. The quantitative panels b) to e) compare all three implementations in terms of correlation-based dependence, mutual-information-based dependence, best-match fluorophore recovery, and a grouped direct same-index GT-versus-output channel-correlation plot. In this 3-channel benchmark, the MATLAB-style workflows preserve fluorophore identity almost perfectly, whereas the source-sink formulation suppresses dependence more strongly but with a noticeable loss in recovery fidelity.

**Fig. 5.**
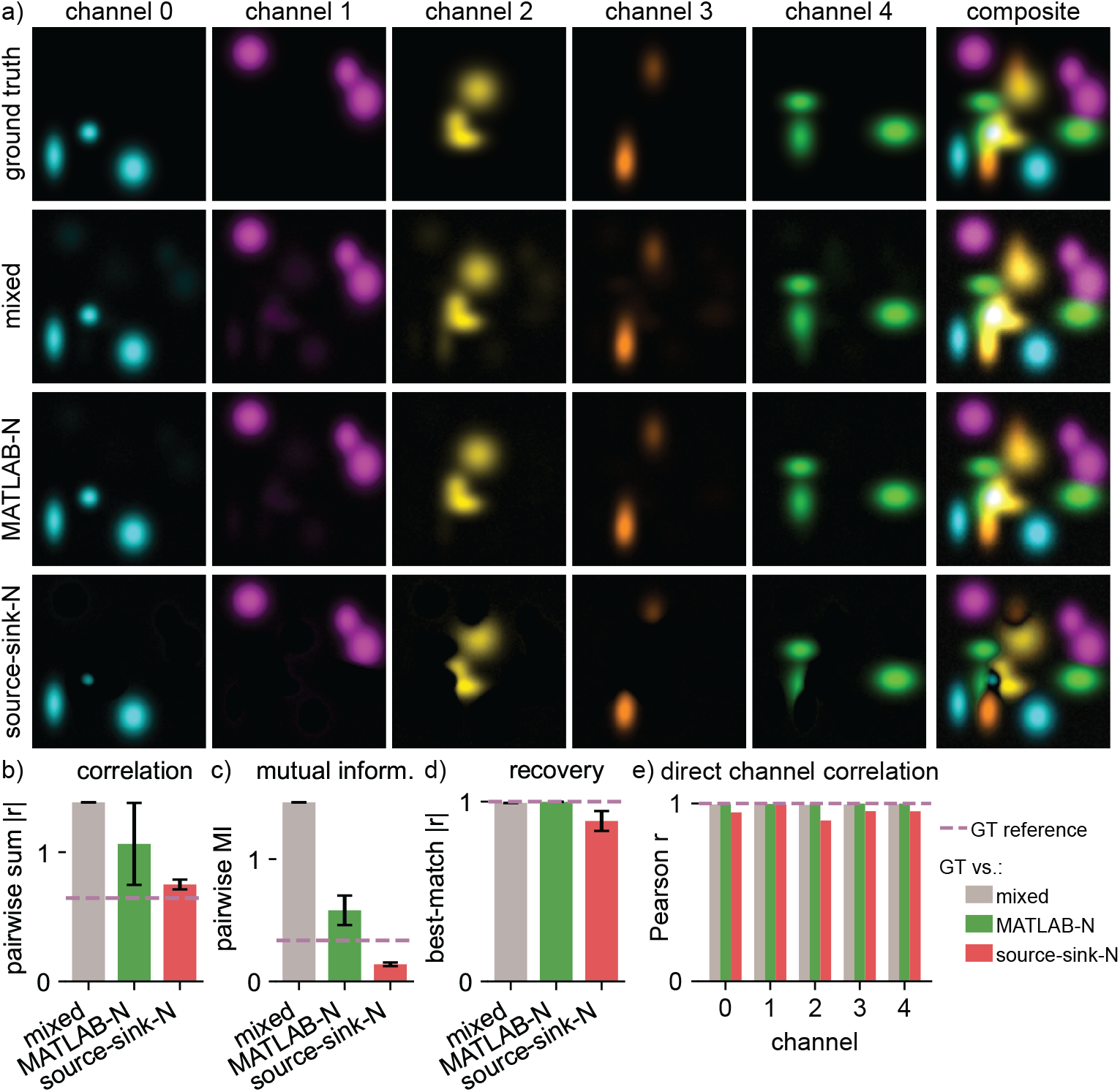
PICASSO-family blind-unmixing benchmark: 5-channel case. Panel a) shows individual panels of the ground-truth fluorophore channels, the spectrally mixed measurements, and the unmixed outputs obtained by the MATLAB-style *N*-channel and source-sink *N*-channel workflows. The quantitative panels b) to e) compare both implementations in terms of correlation-based dependence, mutual-information-based dependence, best-match fluorophore recovery, and a grouped direct same-index GT-versus-output channel-correlation plot. The 5-channel case reproduces the same trade-off seen in the 3-channel benchmark in Fig. 4: MATLAB-style updates are more conservative and preserve fluorophore identity better, whereas the source-sink formulation reduces dependence more aggressively on unrestricted all-to-all mixtures.

The source-sink formulation behaved differently. When applied with an all-to-all source-sink graph on the same synthetic mixtures, source sink n reduced dependence more aggressively, reaching a 3-channel correlation sum of 0.268 ± 0.055 and a mutual-information sum of 0.061 ± 0.025 (Fig. 4b,c). However, this stronger suppression came at a cost in fluorophore fidelity, with the best-match recovery correlation dropping to 0.904 ± 0.062 (Fig. 4d). The same trade-off appeared in the 5-channel benchmark: MATLAB-style *N*-channel unmixing reduced the correlation sum from 1.385 ± 0.365 to 1.064 ± 0.317 and the mutual-information sum from 1.464 ± 0.284 to 0.582 ± 0.121, while maintaining a best-match recovery correlation of 0.9987 ± 0.0002. In contrast, source sink n reduced those dependence scores further to 0.750 ± 0.038 and 0.140 ± 0.015, respectively, but with best-match recovery reduced to 0.892 ± 0.055 (Fig. 5b–d). This behaviour is important to interpret correctly. In these synthetic PICASSO benchmarks, source sink n was intentionally run with an unrestricted all-to-all interaction graph, so that every non-diagonal channel relation was treated as a permissible source-to-sink pathway. That setting is deliberately stressful for the source-sink formulation because it removes the directional prior knowledge on which the method is conceptually based. Under such conditions, each sink is optimized independently to minimize the summed mutual information between the corrected sink and all allowed sources. The easiest route to lower dependence can then become overly aggressive subtraction of shared image structure, rather than recovery of a globally consistent fluorophore decomposition. After positivity clipping, this over-subtraction appears visually as partially hollowed channels or local “holes” in the recovered images. The effect was numerically substantial in the representative benchmark scenes, where many learned off-diagonal source-sink coefficients approached unity and large fractions of sink pixels were driven exactly to zero after correction.

These synthetic all-to-all mixtures therefore highlight an important methodological distinction. The MATLAB-style PICASSO workflows are comparatively conservative and preserve fluorophore identity extremely well, whereas the source-sink formulation acts as a stronger dependence-suppression operator that should ideally be used when the source-sink graph encodes meaningful prior structure rather than unrestricted all-to-all coupling.

### 4.5 Biological example images reproduce the same qualitative trends

In addition to the synthetic benchmarks, we applied the package to real biological microscopy examples and real-data-derived spectral-mixing examples[27]. On the 2-channel biological image derived from the PICASSO figure dataset[29], all five directed linear-unmixing methods reduced residual source-target dependence relative to the raw measurement (Fig. 6). The raw two-channel correlation score of 0.806 decreased to 0.588 for the manually tuned fixed-alpha correction, to 0.544 for mean-ratio estimation, to 0.545 for linear-fit estimation, to 0.551 for correlation minimization, and to 0.655 for mutual-information minimization. The corresponding mutual-information score decreased from 0.156 in the raw image to 0.085 (manual), 0.064 (mean-ratio), 0.065 (linear-fit), 0.068 (correlation minimization), and 0.103 (mutual-information minimization). However, the negative-fraction metric shows that these lower dependence scores were not equally conservative. The fraction of target pixels that would have become negative before clipping was 0.416 for the manual setting, but increased to 0.815 for mean-ratio, 0.811 for linear-fit, and 0.761 for correlation minimization, whereas mutual-information minimization remained much less aggressive at 0.137. Thus, on this experimental example, the methods that most strongly reduced dependence according to correlation and mutual information were also those that relied most heavily on post hoc clipping, while mi min achieved a more conservative subtraction at the cost of leaving more residual cross-channel dependence.

**Fig. 6.**
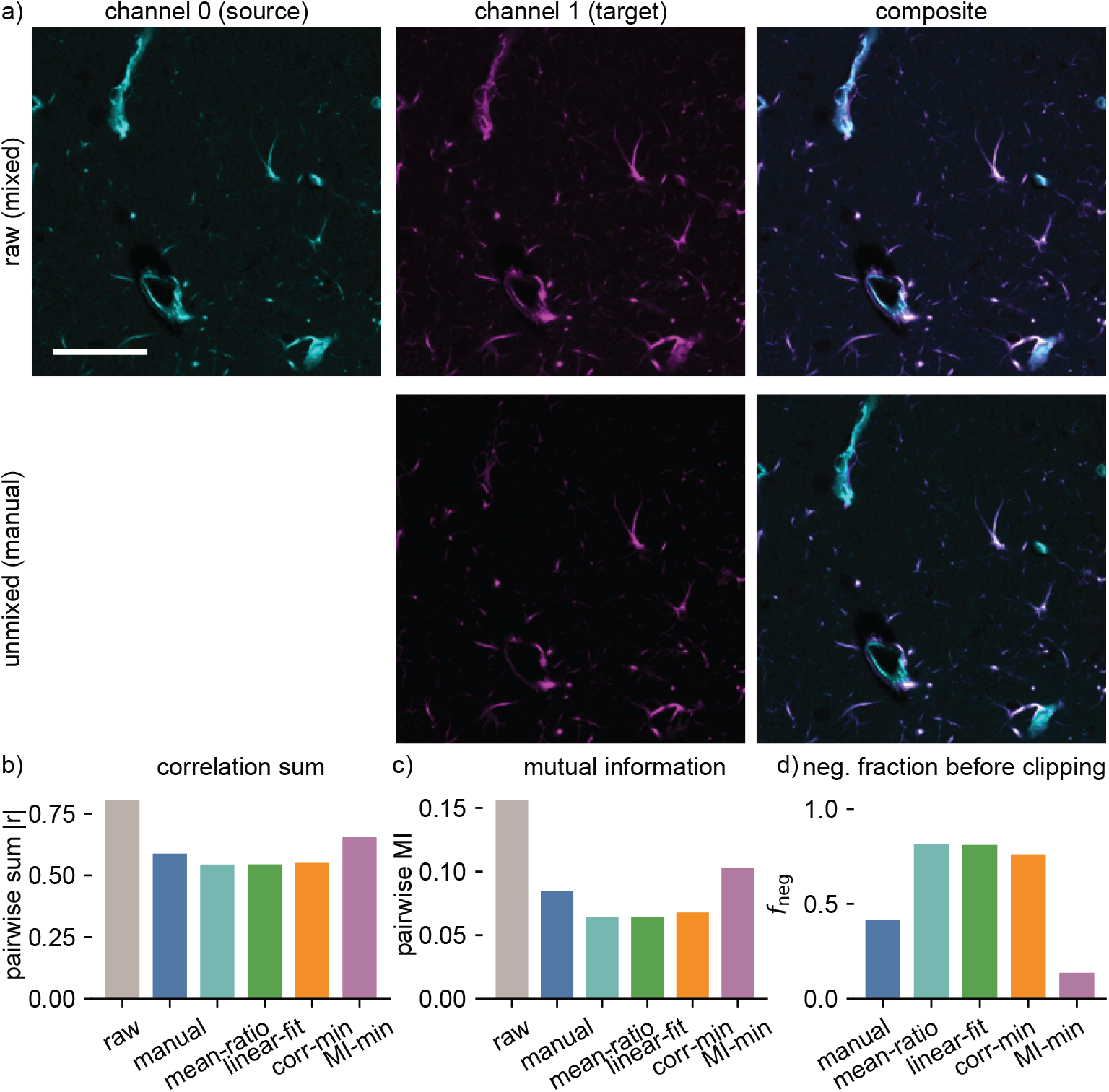
Two-channel biological example. Panel a) shows the raw source channel (cyan), the raw target channel (magenta), and the raw composite in the top row, together with the manually tuned fixed-alpha target-channel result and composite in the bottom row. Panels b) and c) summarize the pairwise sum *|r|* and pairwise mutual information across the raw measurement and all five directed linear-unmixing methods. Panel d) shows the pre-clipping negative fraction *f*_neg_, that is, the fraction of target pixels that would become negative before enforcing non-negativity. For panels b) and c), lower values indicate reduced residual channel coupling after correction; for panel d), higher values indicate more aggressive subtraction and greater reliance on clipping. The scale bar corresponds to 50 *µ*m. The source data originate from the PICASSO figure dataset[29].

On the 5-channel example images from the PICASSO figure dataset, real microscopy source images served as channel-wise ground truth, and measured mixed channels were generated by applying a known fluorophore mixing matrix and additive noise[29]. Panel a) of Fig. 7 shows the ground-truth source-image stack (upper panels), the measured mixed stack (middle panels), and the MATLAB-style *N*-channel result (lower panels). Quantitatively, the measured correlation sum dropped from 4.694 to 2.467 after MATLAB-style *N*-channel unmixing, while the mutual-information sum decreased from 1.406 to 0.308 (Fig. 7b,c). At the same time, mean ground-truth-matched channel recovery increased from 0.566 in the measured stack to 0.799 (Fig. 7d). The direct same-index channel-correlation (Fig. 7e) resolves this recovery pattern at the level of individual channels. Before unmixing, measured-versus-ground-truth correlations were already very high for channels 0 and 4 (*r* ≈ 0.995 and 0.999), but substantially lower for channels 1–3 (*r* ≈ 0.240, 0.475, and 0.121). After MATLAB-style *N*-channel unmixing, channels 1 and 2 improved strongly (*r* ≈ 0.756 and 0.739), channel 3 improved more moderately (*r* ≈ 0.501), and channels 0 and 4 remained near-perfect (*r* ≈ 0.998 and 0.999). The ground-truth stack itself had much lower reference dependence values (0.624 for correlation sum and 0.063 for mutual information), so the result did not reach ideal separation. However, it moved markedly closer to the underlying fluorophore channels.

**Fig. 7.**
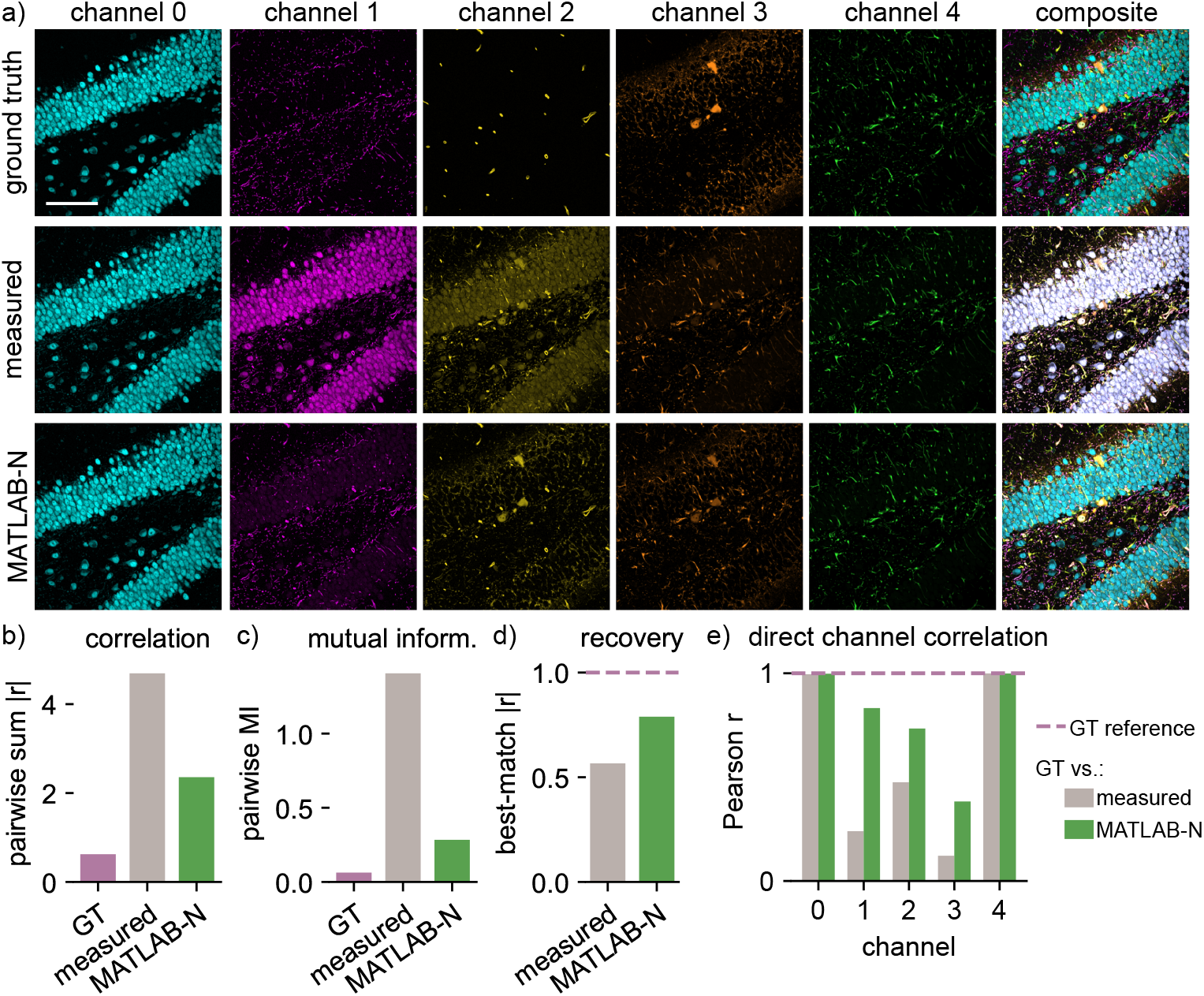
Five-channel real-data-derived simulation example. Panel a) shows the ground-truth source-image stack (top row), the spectrally mixed measured stack (middle row), and the MATLAB-style *N*-channel blind-unmixing output (bottom row). The ground-truth channels are real microscopy images from a mouse brain slice, whereas the spectrally mixed measurements were synthetically generated from these channels using a fluorophore mixing matrix and additive noise in the original PICASSO study. Within each row of panel a), the first five subpanels show individually colorized channels and the last subpanel shows the composite. Panels b) to e) summarize pairwise sum *|r|*, pairwise mutual-information sum, ground-truth-matched recovery, and direct same-index GT-versus-output channel correlations. For panels b) and c), lower values indicate reduced residual channel coupling after unmixing. For panels d) and e), higher values indicate better correspondence to the underlying fluorophore channels. The scale bar corresponds to 50 mm (5 cm). The source data originate from the PICASSO figure dataset[29].

The incomplete recovery in this 5-channel example is plausible and informative rather than surprising. In contrast to the near-ideal synthetic benchmarks, this real-data-derived simulation contains more heterogeneous channel structure, stronger overlap patterns, and a higher-dimensional parameter space in which the sequential MATLAB-style pairwise updates become a progressively more approximate surrogate for the true latent decomposition. Put differently, the workflow still drives the measured channels substantially toward the ground truth, but some residual ambiguity remains because the blind-unmixing objective is optimized only indirectly through pairwise dependence-reduction steps rather than through access to the true mixing matrix. The example therefore illustrates both the practical usefulness and the natural limits of the generalized MATLAB-style *N*-channel formulation.

This example also clarifies an important practical trade-off in higher-channel settings: optimizing parameters toward lower residual dependence alone does not necessarily maximize correspondence to the underlying fluorophore channels. In this five-channel example, a finer parameter setting improved ground-truth recovery even though it did not drive the dependence scores down as far as a more subtractive alternative would have done. In other words, stronger decorrelation is not automatically the same as better fluorophore reconstruction once the problem becomes sufficiently high-dimensional.

### 4.6 Directional source-sink unmixing reduces dependence while retaining sink-channel structure

The all-to-all synthetic benchmarks above intentionally stress-test the source-sink formulation outside its most natural use case. A more realistic directional example is the real-data-derived GFAP sink LMNB1 source image from the PICASSO figure dataset[29], in which channel 1 is a plausible contamination source for channel 0. The image panels in Fig. 8a compare the raw source-sink pair with the MATLAB-style *N*-channel and source sink n results. On this image, the raw pairwise correlation score of 0.217 dropped to 0.0779 after MATLAB-style *N*-channel blind unmixing and further to 0.0318 after source sink n correction (Fig. 8b). The corresponding mutual-information score decreased from 0.0886 to 0.0317 for MATLAB-style *N*-channel unmixing and to 0.0369 for source sink n (Fig. 8c).

**Fig. 8.**
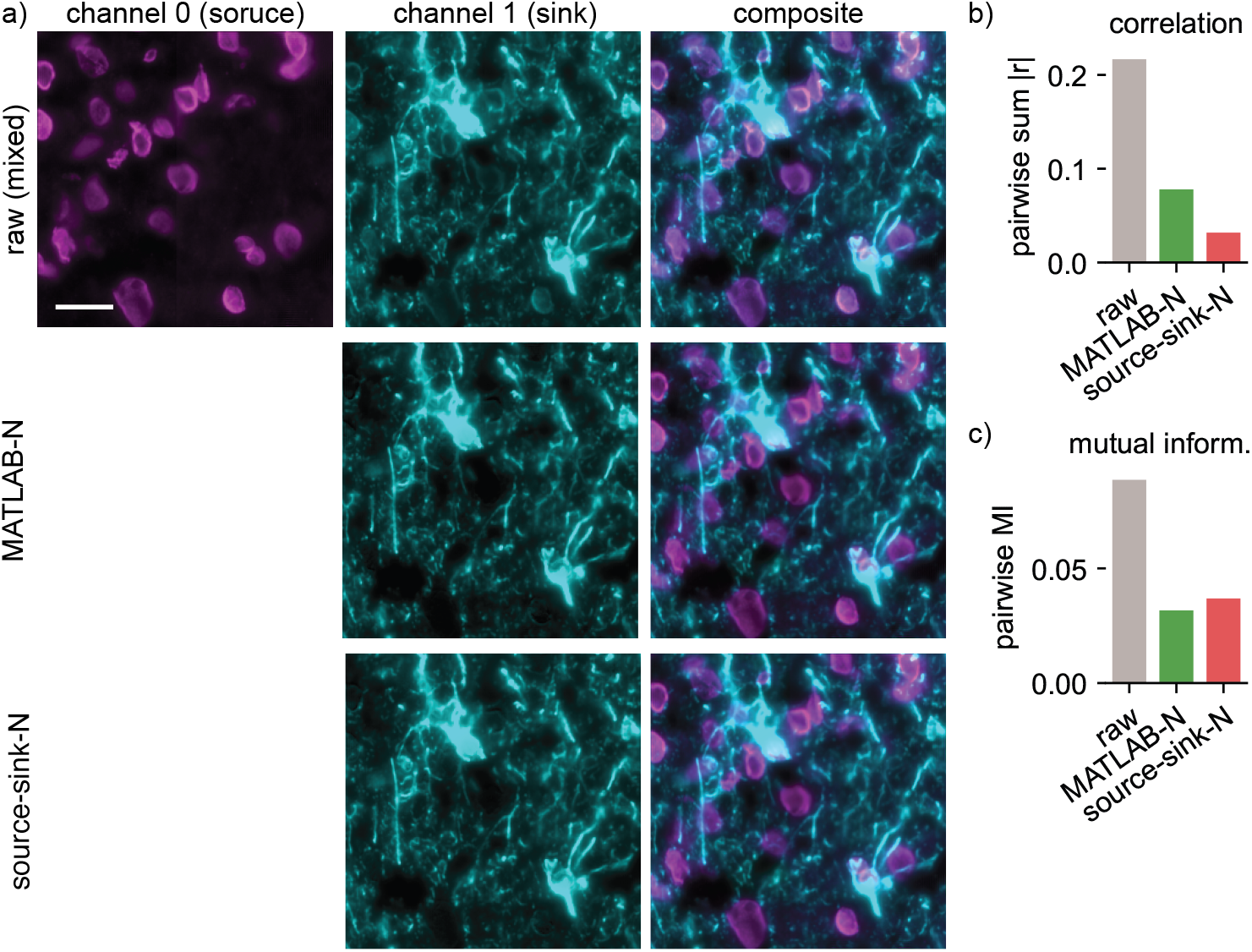
Directional source-sink example from a real microscopy dataset. Panel a) shows the raw source channel (magenta), the raw sink channel (cyan), and the raw composite in the top row. The middle row of panel a) shows the MATLAB-style *N*-channel unmixed sink together with its composite using the raw source channel, and the bottom row shows the corresponding source sink n sink and composite. Panels b) and c) summarize the pairwise sum *|r|* and pairwise mutual information. In both panels, lower values indicate reduced residual channel coupling after unmixing. The scale bar corresponds to 50 *µ*m. The example illustrates the intended use case of the source-sink formulation: explicit directional priors can reduce cross-channel coupling without forcing an unrestricted all-to-all blind-unmixing model. The source data originate from the PICASSO figure dataset[29].

These scores should again be interpreted as within-image comparisons rather than as universal thresholds. Nevertheless, the example is informative because it complements the all-to-all synthetic mixtures: the MATLAB-style workflow suppresses dependence strongly but also over-removes the designated sink channel visually, whereas the directional source-sink workflow retains more sink-channel structure while still reducing cross-channel coupling substantially. This is the regime in which source sink n is most conceptually appropriate.

## 5 Discussion

*spectral-unmixing* turns bleed-through correction from a fragmented, often laboratory-specific processing step into a reproducible and extensible analysis workflow for multidimensional microscopy data by bringing together several linear spectral-unmixing workflows that are often treated separately in practice: directed two-channel correction with multiple alpha estimators, optional bidirectional correction via a proper 2 × 2 inverse model, and multi-channel PICASSO-family blind unmixing. By placing these workflows under one API and pairing that API with canonical multidimensional stack handling and machine-readable sidecar reports, the package makes routine spectral-unmixing analyses easier to reproduce, audit, and share.

The synthetic benchmarks show that relatively simple estimators, such as the mean-ratio and linear-fit approaches, performed very well in the two-channel correction setting. This is useful because many practical microscopy workflows do not need the most sophisticated possible optimizer. Instead, they need a transparent estimator that can be reasoned about scientifically and reproduced later. The time-varying benchmark further highlights that the package’s explicit alpha modes are not a cosmetic design choice. When bleed-through changes across time, per-time-point estimation can recover that drift far more faithfully than a single reference-time-point estimate.

Within the blind-unmixing framework, three workflows serve complementary purposes: a 3-channel path, an *N*-channel path, and a source-sink *N*-channel path. The 3-channel path provides a close Python port of the iterative logic used in the original PICASSO MATLAB workflow[4]. The generalized MATLAB-style *N*-channel path extends the same pairwise-update idea to larger channel sets, whereas the source-sink formulation allows users to encode prior knowledge about plausible contamination directions. The benchmark results highlight that these workflows are not interchangeable. The MATLAB-style workflows behaved conservatively on synthetic all-to-all mixtures and preserved fluorophore identity extremely well, while the source-sink formulation acted as a stronger dependence-suppression operator whose behaviour depends more directly on the chosen interaction graph. In particular, the “hollowed” appearance of some source-sink benchmark images reflects over-subtraction under an unrestricted all-to-all interaction graph rather than a numerical instability. When an explicitly directional subtraction model is applied without directional priors, every channel is allowed to explain every other channel. Independent per-sink optimization can then remove shared structure so aggressively that positivity clipping leaves visible voids in the recovered channels. The five-channel real-data-derived simulation from the PICASSO figure dataset is informative in the same regard: with tuned settings, the MATLAB-style *N*-channel path improved channel-wise correspondence to ground truth substantially, but did not produce the lowest possible dependence scores. In practical use, the most suitable blind-unmixing path therefore depends on the imaging design, the number of measured channels, and whether the immediate priority is stronger dependence suppression, better fluorophore recovery, or a compromise between both.

Several limitations are worth stating clearly. First, *spectral-unmixing* is built around linear mixing assumptions. This is appropriate for many microscopy bleed-through problems, but it does not address nonlinear detector behaviour or more complex optical interactions. Second, data-driven alpha estimation from experimental images can be biased when genuine target signal overlaps strongly with source signal. For that reason, a manually provided alpha measured on a single-label control remains scientifically preferable whenever such a control is available. Third, the source-sink blind-unmixing workflow benefits from user knowledge: if the chosen source-sink graph is biologically implausible, the resulting correction may also be implausible. The synthetic all-to-all PICASSO benchmark results make the same point from the opposite direction: when the source-sink workflow is used without meaningful directional priors, it can suppress dependence strongly but also deviate further from the underlying fluorophore decomposition than the more conservative MATLAB-style workflows, including the visually conspicuous formation of clipped low-intensity voids after over-subtraction. Fourth, beyond three channels the MATLAB-style matlab n workflow should be interpreted as a transparent and useful package-level generalization of the released PICASSO MATLAB logic, not as an original high-channel reference implementation. In higher-dimensional examples this means that the method can move the data substantially toward the latent fluorophore channels without necessarily reaching a fully resolved decomposition, because the optimization proceeds through sequential pairwise updates rather than direct recovery of the true global mixing matrix. The real-data-derived examples suggest that this generalization can still improve recovery substantially, but also that parameter tuning and metric choice matter more strongly in the higher-channel regime.

These limitations do not weaken the core motivation for the framework. Rather, they define a realistic and useful software scope: *spectral-unmixing* is a platform for reproducible linear unmixing workflows, not a universal substitute for experimental controls or physical calibration. Within that scope, its modular structure is a strength. New alpha estimators, new blind-unmixing routines, and new pre-or post-processing helpers can be added incrementally without reworking the rest of the code base. This makes the package suitable not only for immediate use but also for community-driven extension and long-term maintenance.

## 6 Conclusions

*spectral-unmixing* addresses a practical and reproducibility-oriented gap in microscopy image analysis: it provides a single Python package for directed, bidirectional, and blind spectral-unmixing workflows in multidimensional microscopy stacks. It standardizes how common linear bleed-through correction workflows are expressed, executed, documented, and shared. By combining canonical multidimensional stack handling, explicit mathematical models, multiple coefficient-estimation strategies, and sidecar reports, the package turns spectral unmixing from an often manual or laboratory-specific procedure into a reproducible, shareable, and FAIR-oriented research-software workflow.

## 7 Availability and requirements

- **Project name:** *spectral-unmixing*.
- **Project home page:** https://github.com/FabrizioMusacchio/Spectral_Unmixing.
- **Archived version:** The software is archived on Zenodo[25].
- **Operating system:** Platform independent; tested primarily on macOS and Linux through continuous integration.
- **Programming language:** Python 3.12 or newer.
- **Key dependencies:** NumPy, SciPy, scikit-image, pystackreg, matplotlib, napari for optional visualization, and OMIO for microscopy image I/O.
- **Documentation:** https://spectral-unmixing.readthedocs.io/en/latest/.
- **Example data:** Companion example datasets are archived separately on Zenodo[27].
- **License:** GNU General Public License v3.0 or later (GPL-3.0-or-later).
- **Restrictions to use by non-academics:** None beyond the terms of the open-source license.

## List of abbreviations

*API*: Application programming interface.
*FAIR*: Findable, accessible, interoperable, and reusable.
*GPL*: GNU General Public License.
*I/O*: Input/output.
*MI*: Mutual information.
*NRMSE*: Normalized root mean squared error.
*OME*: Open Microscopy Environment.
*OMIO*: Microscopy image input/output library used for reading and writing microscopy stacks.
*PICASSO*: Blind-unmixing method family introduced for ultra-multiplexed fluorescence imaging without reference spectra.
*TZCYX*: Canonical stack axis order: time, z, channel, y, x.

## Declarations

### Ethics approval and consent to participate

Not applicable. This manuscript does not report studies involving human participants, human data, human tissue, animals, or client-owned animals.

### Consent for publication

Not applicable. The manuscript does not contain any individual person’s data.

### Availability of data and materials

The companion example datasets used in the tutorials and qualitative figures are archived on Zenodo[27]. The source datasets from which the included biological and real-data-derived examples were derived are cited in the corresponding figure legends and bibliography. The *spectral-unmixing* package is available on GitHub at https://github.com/FabrizioMusacchio/Spectral_Unmixing and archived on Zenodo[25]. The documentation is available at https://spectral-unmixing.readthedocs.io/en/latest/. The scripts used to generate the synthetic quantitative benchmarks and manuscript figures are included in the repository under additional scripts/.

### Competing interests

The authors declare that they have no competing interests.

### Funding

MF received funding from the European Research Council (ERC; MicroSynCom 865618) and the German Research Foundation (DFG; SFB1089 C01, B06; SPP2395).

### Authors’ contributions

F.M. conceived the package, implemented the software, designed and performed the benchmark analyses, prepared the figures, and wrote the manuscript. M.F. supervised the project and provided institutional and scientific oversight. Both authors read and approved the final manuscript.

## Acknowledgements.

We thank colleagues and collaborators for discussions around microscopy I/O, interoperable workflow design, and reproducible bioimage analysis. We also thank the Fiji/ImageJ and broader open-source bioimage-analysis communities for building the software ecosystem on which this work relies, and the authors of the original PICASSO publication for releasing both the method and example data that enabled transparent benchmarking of blind-unmixing workflows.

## Use of AI-assisted language tools

The authors used OpenAI ChatGPT 5.5 for language correction and proofreading only. The authors subsequently reviewed and edited the manuscript and take full responsibility for the content of the article.

## Notes

### Competing Interest Statement

The authors have declared no competing interest.

### Summary of Updates

We have substantially revised the manuscript structure. The Background and Implementation sections were revised to clarify the software architecture, image input and axis handling, modular unmixing design, and reproducibility features. The abstract and conclusions were sharpened to better summarize the scope and main findings. Figure 1 was replaced with an updated software architecture schematic. The Results and Discussion sections were revised to clarify interpretation of the directed, bidirectional, and PICASSO family benchmark results, including the role of source sink priors and residual dependence metrics. Figure captions and metric descriptions were updated for clarity. Declarations, availability statements, citations, and wording related to software and data availability were also updated.

